# Monocytes from Older Adults Maintain Capacity for Metabolic Compensation during Glucose Deprivation and Lipopolysaccharide Stimulation

**DOI:** 10.1101/633354

**Authors:** Johnathan R. Yarbro, Brandt D. Pence

**Affiliations:** School of Health Studies, University of Memphis, 38152, USA; Center for Nutraceutical and Dietary Supplement Research, University of Memphis, Memphis, TN 38152, USA

## Abstract

Inflammaging is the chronic low-grade inflammation that occurs with age that contributes to the pathology of age-related diseases. Monocytes are innate immune cells that become dysregulated with age and which can contribute to inflammaging. Metabolism plays a key role in determining immune cell functions, with anti-inflammatory cells primarily relying on fatty acid oxidation and pro-inflammatory cells primarily relying on glycolysis. It was recently shown that lipopolysaccharide (LPS)-stimulated monocytes can compensate for a lack of glucose by utilizing fatty acid oxidation. Given that mitochondrial function decreases with age, we hypothesized that monocytes taken from aged individuals would have an impaired ability to upregulate oxidative metabolism along with impaired effector functions. Aging did not impair LPS-induced oxygen consumption rate during glucose deprivation as measured on a Seahorse XFp system. Additionally, aged monocytes maintained inflammatory gene expression responses and phagocytic capacity during LPS stimulation in the absence of glucose. In conclusion, aged monocytes maintain effector and metabolic functions during glucose deprivation, at least in an ex vivo context.

---

The number of Americans over the age of 65 is expected to increase by approximately 80 million people by 2040^1^. This change will have significant effects on the economy, healthcare, and society in general. Currently, over two-thirds of the healthcare budget is spent on managing chronic diseases of the elderly^2^. Aging is the highest risk factor for the majority of chronic diseases - including cardiovascular disease, diabetes, arthritis, and cancer^3^. While lifespan continues to rise, healthspan (the length of time someone is healthy) has increased more slowly, and Americans are living longer with impaired health and disabilities^4^. Interventions are needed to improve the health and quality of life of the aging population, and studies show that the compression of morbidity is possible with lifestyle changes, pharmaceuticals, and continuous medical advances^3, 5^.

Aging is associated with chronic, low-level, systemic inflammation (termed inflammaging) that contributes to most, if not all, age-related diseases^6^. Older adults have higher serum levels of several pro-inflammatory cytokines/proteins such as IL-6, IL-1*β*, C-reactive protein (CRP), and TNF*α*^7^. Elevated levels of these molecules in circulation, most notably IL-6, are correlated with an increased risk of morbidity and mortality in elderly populations^8^. Furthermore, they are associated with sarcopenia, malnutrition, arthritis, atherosclerosis, cognitive decline, and other diseases of aging^9^. There is no consensus on the causes of inflammaging, though it is likely due to a host of factors that become dysregulated with age. These “hallmarks of aging” include reduced autophagy and mitophagy, accumulation of DNA and mtDNA damage leading to genomic instability, epigenetic changes, telomere shortening, cellular and immune senescence, dysbiosis, chronic antigenic stress, diminished proteostasis, altered metabolic signaling, stem cell exhaustion, increased cellular garbage, and mitochondrial dysfunction^10^.

Monocytes are circulating mononuclear phagocytes of the innate immune system that have a diverse set of functions and play an essential role in the defense against a variety of pathogens^11^. Although aging is multi-factorial, monocytes may play an essential role in aging pathology. Their heterogenous and highly adaptable nature, ability to respond to pathogens and cellular garbage, communication with the adaptive immune system, and numerous defects with age make them a key contributor to inflammaging. Monocytes from older individuals have impaired migration and phagocytosis, diminished ability to activate the adaptive immune system, altered receptor expression, cytokine production, and subset proportions^12, 13^.

In response to activation, immune cells regularly change their functions dramatically. For example, monocytes activated by lipopolysaccharide (LPS) undergo vigorous cellular growth and an increased demand for the production of proteins, lipids, cytokines, and reactive oxygen species (ROS)^14, 15^. This increased demand for biomolecules is associated with upregulation of glycolysis and downregulation of oxidative phosphorylation (OXPHOS), similar to the Warburg effect in cancer cells^16^. A shift toward aerobic glycolysis is associated with pro-inflammatory phenotypes in immune cells such as dendritic cells activated by LPS^17^, M1 macrophages, and effector T cells^18^. Oxidative metabolism (with limited rates of glycolysis) is associated with anti-inflammatory phenotypes as in quiescent memory T-cells, regulatory T-cells, and M2 macrophages^18^. The metabolic inflexibility of an immune cell can lead to increased inflammation. Macrophages are able to convert from one form to another and can switch their metabolism during inflammation from relying on glycolysis in the M1 state to relying on OXPHOS in the M2 state^19^. Inhibition of OXPHOS in macrophages inhibits the expression of the M2 anti-inflammatory phenotype^20^. Therefore, diminished mitochondrial function is presumed to affect the function and phenotype of certain immune cells.

Aging has been shown to cause decreased mitochondrial function with an estimated decline of 8% in ATP producing capacity per decade^21^. This decrease in function is thought to be largely caused by mitochondrial DNA (mtDNA) mutations, which occur at an estimated rate of 15x that of the nuclear genome^21^. Since mitochondrial dysfunction is impaired it is reasonable to assume that many cell types in older individuals must produce more energy through non-oxidative metabolism. Aged mice have been shown to have increased lactate and reduced glycolytic intermediates in muscle and liver tissue which suggest an increased reliance on anaerobic glycolysis^22^. Whether or not mitochondrial dysregulation is a primary cause of age-related monocyte dysfunction has yet to be determined, though we’ve recently provided evidence that aging impairs maximal and spare mitochondrial respiratory capacity in classical monocytes isolated from humans^13^.

Raulien and colleagues recently showed that in the deprivation of glucose (as is often the case in tissues with inflammation)^23, 24^, LPS activated monocytes are able to switch from aerobic glycolysis to OXPHOS while maintaining their effector functions^14^. This switch is regulated by AMP-activated protein kinase (AMPK) which stimulates catabolic pathways including fatty acid oxidation, autophagy, and mitochondrial biogenesis^14^. Since we have previously demonstrated mitochondrial dysfunction in monocytes^13^, we hypothesized that aging would result in a reduced ability to upregulate oxidative metabolism when glucose is unavailable following LPS-stimulation and that this would be associated with reduced inflammation and phagocytic capacity.

## Results

### Data availability

The datasets and analytical scripts supporting the conclusions of this article are available in the FigShare repository^25^.

### Subject characteristics

Subject demographics and anthropometric data are shown in Table 1. There was a total of 9 subjects in the aged group and 11 subjects in the young group. Besides age, the two groups did not differ significantly on other demographic or anthropometric data. All subjects are the same individuals as those reported in our previous paper as cohort 2^26^.

**Table 1.** Demographic and anthropometric characteristics of subjects. Abbreviations: yr, year; cm, centimeters; kg, kilograms; BMI, body mass index; N, number of subjects

| Characteristics | Aged (n=9) | Young (n=11) | Probability |
| --- | --- | --- | --- |
| Age, yr. (range) | 66.6 ± 1.4 (60-72) | 27.5 ± 1.3 (20-34) | p < 0.001 |
| Height, cm (range) | 173.8 ± 3.0 (163.0-191.0) | 169.8 ± 3.2 (155.0-191.0) | p = 0.375 |
| Weight, kg (range) | 77.3 ± 4.0 (54.4-99.8) | 74.8 ± 7.5 (43.4-133.8) | p = 0.789 |
| BMI, kg/m<sup>2</sup> (range) | 25.6 ± 1.2 (18.8-31.6) | 25.7 ± 2.0 (17.4-36.7) | p = 0.980 |
| Female, N (%) | 4 (44) | 4 (36) | p = 0.714 |
| White, N (%) | 7 (78) | 4 (36) |  |
| Black, N (%) | 2 (22) | 2 (18) | Race: p = 0.059 |
| Asian, N (%) | 0 (0) | 5 (45) |  |

### Metabolic flexibility assay, cytokine expression, and phagocytic capacity

Aging had no significant effects on any of the calculated measures for metabolic flexibility, cytokine expression, or phagocytic capacity in CD14+CD16- monocytes. Oxygen consumption rate (OCR) response to LPS in glucose-deprived monocytes was slightly higher across all timepoints in the aged group, though this difference was not significant (Fig. 1a). Extracellular acidification rate (ECAR) response to LPS in glucose-deprived monocytes also showed no difference between groups (Fig. 1b). Aging had no effect on any of the calculated oxidative metabolic parameters (Fig. 1c & d) including maximum OCR (p=0.3312), minimum OCR (p=0.3702), the difference between them (p=0.2206), and OCR kinetics as measured by area under the curve (AUC) (p=0.3312). Aging also had no significant effects on measures of cytokine expression in monocytes in response to LPS (Fig. 1e) for IL-10 (p=0.5675), IL-6 (p=0.8421), TNF*α* (p =0.9048), or IL-1*β* (p=0.7802).

**Figure 1.**
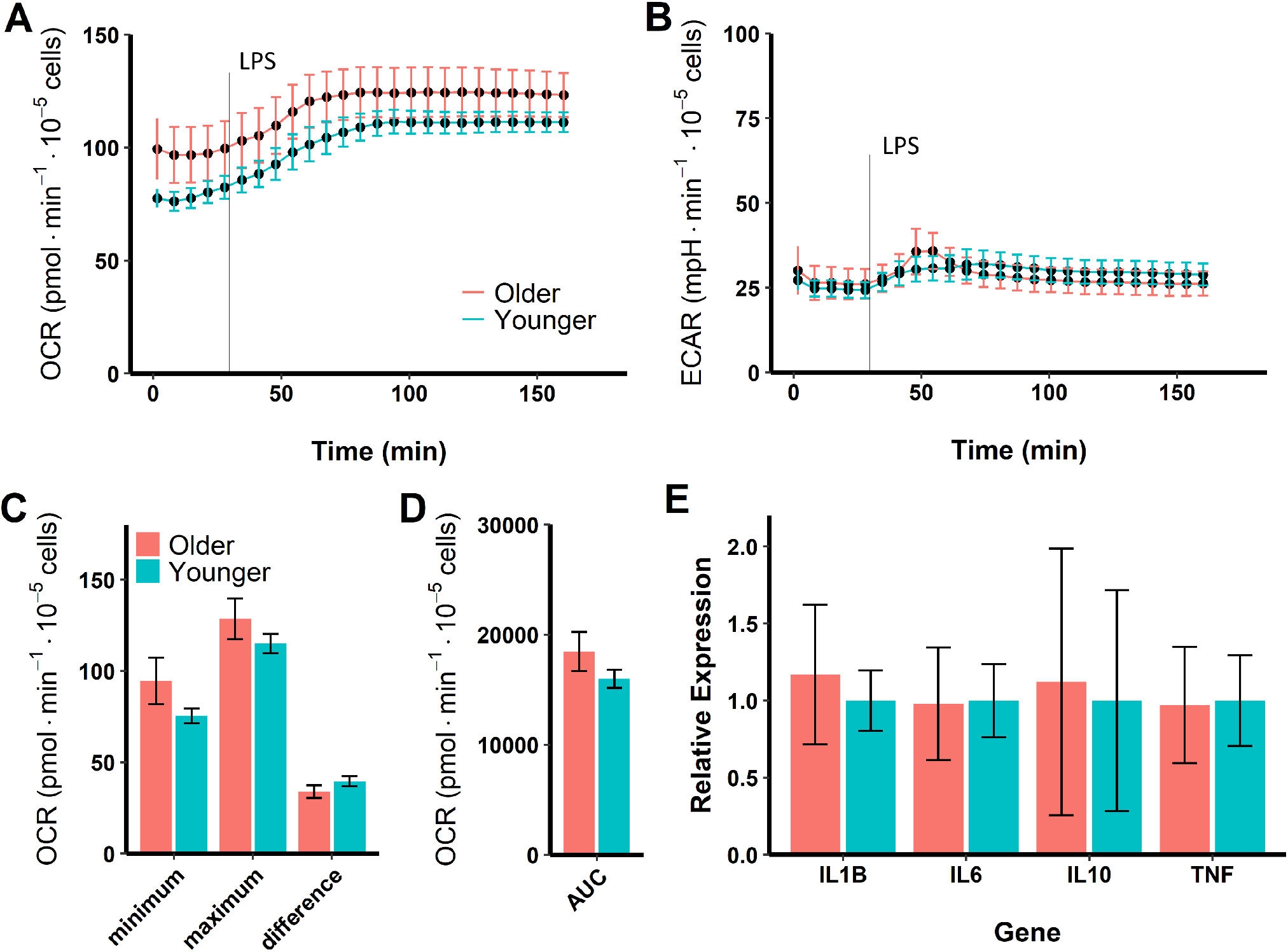
Metabolic responses to LPS in glucose-deprived CD14+CD16- monocytes. **A**) Oxygen consumption response to LPS in older and younger subjects. **B**) Glycolytic response to LPS. **C**) Calculated values for respiratory parameters. **D**) Kinetic OCR response (AUC: area under the curve). **E**) Cytokine gene expression.

Phagocytic capacity, as measured by percentage in the positive gate, was tested in glucose-deprived classical monocytes (Fig. 2). An example of the gating strategies used to isolate classical monocytes (Fig. 2a) and to calculate percentage positive for beads (Fig.2b) are shown. Aging had no effect on the mean fluorescence intensity (MFI) in LPS-stimulated (p=0.748, Fig. 2c) or unstimulated (p=0.1396, Fig. 2c) glucose-deprived classical monocytes. Similarly, aging had no effect on percentage of cells in the positive gate (Fig. 2d) in LPS-stimulated (p=0.2340) or unstimulated (p = 0.6738) glucose-deprived classical monocytes. There was a significant main effect whereby LPS stimulation increased MFI compared to unstimulated (p=0.00301, not shown). Likewise, LPS stimulation caused a near significant increase in percentage of cells in the positive gate compared to unstimulated (p=0.0531, not shown).

**Figure 2.**
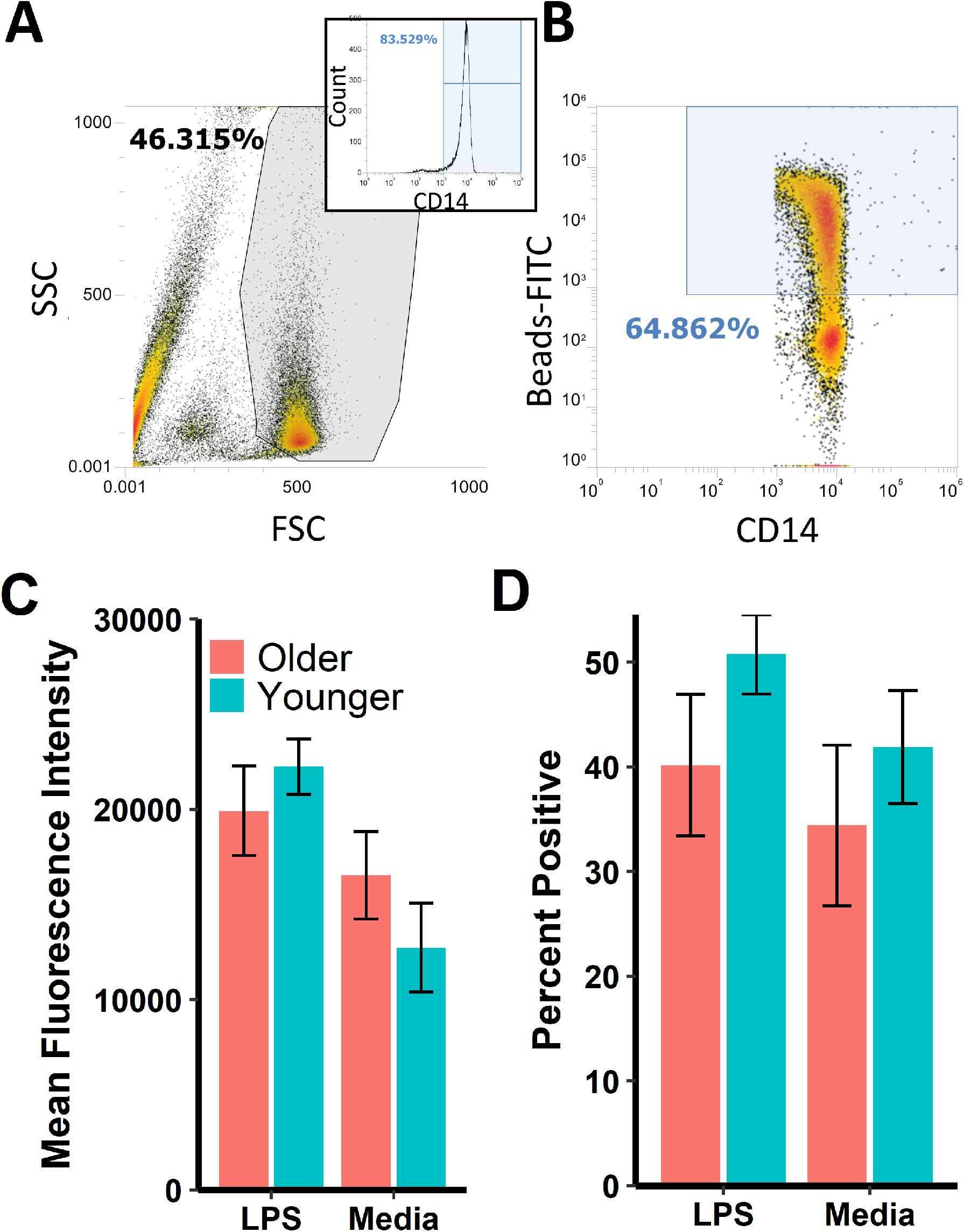
Phagocytic activity in glucose-deprived CD14+CD16- monocytes. **A**) Example of Monocyte Gating Strategy. Top right graph shows percentage of CD14+ cells within gate. **B**) Example of percentage positive for beads phagocytosed by CD14+ cells. **C**) Mean fluorescence intensity in LPS stimulated, or media only CD14+ monocytes in older and younger subjects. **D**) Percentage positive for beads by LPS stimulated, or media only CD14+ monocytes in older and younger subjects.

## Discussion

Monocytes switch from oxidative phosphorylation to aerobic glycolysis when activated by LPS to carry out their effector functions, which is dependent on the availability of glucose and glutamine. Monocytes routinely experience a variety of different microenvironments and must be metabolically flexible to retain their functions. In circulation glucose is readily available, but in areas with inflammation glucose is often drastically reduced, and monocytes must compensate through catabolic processes such as fatty acid oxidation and autophagy^14^. AMPK is the critical regulator orchestrating this metabolic switch, which provides fatty acids to the mitochondria through downstream effects^19^. Decreased mitochondrial function is known to occur in many cell types with age^21^, and we have previously provided evidence for reduced mitochondrial function in classical monocytes with aging– namely that aging impairs mitochondrial maximal respiration and spare capacity^13^.

We hypothesized that classical monocytes from aged individuals would have reduced ability to upregulate oxidative metabolism when glucose is unavailable due to decreased mitochondrial function, and that this inflexibility would impair their inflammatory responses and hamper their ability to perform phagocytosis. In this study, we show that aging has no effect on the ability of glucose-deprived classical monocytes to upregulate oxidative metabolism, and has no effect on the expression of IL-10, IL-6, TNF*α*, or IL-1*β* during acute LPS activation ex vivo. Additionally, we found aging has no effect on ex vivo phagocytic capacity in classical monocytes, regardless of the availability of glucose or whether they were activated by LPS. The differences in mitochondrial respiration in aged monocytes that we previously reported^13^ were only under maximizing conditions (FCCP stimulation), and thus normal conditions are likely not severe enough to cause any noticeable impairments in mitochondrial metabolism.

While there is an ever-growing body of evidence supporting age-related changes in monocyte and macrophage functions, it has not been clear whether this is due to intrinsic changes in the monocytes themselves, or due to changes in circulating factors that alter gene expression. This study gives more credibility to the assumption that classical monocytes are not experiencing intrinsic changes to a degree which affect their functions. Aging may instead result in reductions in monocyte effector functions due to alterations in the in vivo environment, including exposure to inflammatory cytokines, damage-associated molecular patterns, and bacterial products which are increased in the circulation in older adults^8^.

There are several other potential reasons why we observed no differences in this study, which is likely not translatable to monocyte function in vivo. We previously corroborated evidence that alterations in monocyte subset proportions occur with age^13^. CD14+CD16- classical monocytes are reduced, while CD14+CD16+ intermediate and CD14dimCD16+ nonclassical monocytes are increased with age. Our monocyte isolation technique removes CD16+ monocytes and thus only included the classical monocyte subset. This was done to prevent biased results, as CD16+ monocytes have generally shown to be more pro-inflammatory and are more prone to senescence^27^. Furthermore, monocyte subsets have been shown to display varying responses to LPS^28^. Isolating all monocyte subsets therefore may have yielded different results, and it would be interesting to see a similar experiment with respect to metabolic flexibility performed with CD16+ monocytes only, to see if there are intrinsic changes in CD16+ monocytes with age.

Monocytes also have varying responses to different types of pattern recognition receptors (PRR). Activation by Pam3CysSK4, a TLR2 agonist, shows significant differences in TCA cycle, OXPHOS, and lipid metabolism compared to activation by LPS in glucose-fueled monocytes^29^. Therefore, activation of PRRs other than TLR4 may have given different results.

In summary, we showed that aging has no effect on the ability of ex vivo LPS-stimulated classical monocytes to compensate for a lack of glucose by upregulating oxidative metabolism. Furthermore, aging has no effect on cytokine expression in ex vivo LPS-stimulated glucose-deprived classical monocytes for IL-10, IL-6, TNF*α*, or IL-1*β*. Aging also had no effect on the phagocytic ability of classical monocytes under various conditions.

## Methods

### Participants

Males and females between the ages of 18-35 were recruited for the young group, which consisted of 11 subjects total. Males and females between the ages of 60-80 were recruited for the aged group, which consisted of 9 subjects total. Demographic and Anthropometric data of the subjects can be seen in Table 1. All subjects were recruited from the surrounding Memphis area via word-of-mouth, email, or flyers. Exclusion criteria included subjects who had diagnosed conditions that can affect metabolic or immune function. This includes obesity (BMI >30), cardiovascular disease, diabetes, hypertension, chronic fatigue syndrome, mitochondrial diseases, autoimmune diseases, etc. All subjects completed a questionnaire to determine eligibility, which asked for health history, list of medications and supplements, major illnesses or hospitalizations within the last two years, and exercise type/frequency. Subjects visited the lab in a fasted state, up to 6 times, and had 8-16 mL of blood taken by venipuncture into 1-2 EDTA vacutainer tubes per visit for monocyte isolation.

### Monocyte Isolation and Metabolic Flexibility Assay

CD14+ monocytes were isolated from whole blood via magnetic sorting by negative selection using Stemcell Technologies’ EasySep Direct Human Monocyte Isolation Kit. CD16+ monocytes were excluded to prevent bias as monocyte subset proportions change with age^13^. Therefore, in this study we only looked at classical monocytes. After isolation the monocytes were counted using EMD Millipore’s Scepter 2.0 cell counter for use in downstream assays. The Agilent Seahorse XFp Analyzer was used to test the metabolic flexibility of the isolated classical monocytes. The Seahorse analyzer measures the oxygen consumption rate (OCR) and extra-cellular acidification rate (ECAR) of the sample. OCR is an indicator of OXPHOS and ECAR is an indicator of glycolysis. Monocytes were seeded into the Seahorse plates at 1.5 x 10^5^ cells per well (B-G) along with XF media (DMEM, pyruvate, glutamine), and either glucose or no glucose. 20*μ*L of 100ng/mL LPS was added to the injection port of three wells containing glucose (B-D), three wells without glucose (E-G), and two wells (A, H) consisting of no cells, but XF media, which are used as blanks. The Seahorse plates were incubated for 60 minutes at 37°C in a non-CO2 incubator to de-gas the plate. The Seahorse machine was then run for 160 minutes to measure the acute response in OCR and ECAR of the monocytes activated by LPS in media with or without glucose. This data was used to compare the metabolic flexibility of isolated monocytes between the aged and young groups. Wells B-G of the Seahorse plate were imaged using a microscope at 10x magnification to confirm cell adherence and for use in cell number normalization when doing data analysis. After completion, wells B-G received 100*μ*g of TRIzol, pooled by group (+ or - glucose) and were stored in a −80°C freezer for later RNA quantification using Real-Time polymerase chain reaction (qPCR).

### Phagocytosis Assay

To test potential differences in monocyte effector function between young and aged CD14+ CD16- monocytes, a phagocytosis assay was used in the presence and absence of glucose. The assay uses latex beads coated with fluorescently labeled IgG to quantify phagocytosis in vitro and was measured using the Attune Nxt flow cytometer. Monocytes were isolated as above and added to 2mL tubes at a concentration of 5 x 10^5^ cells. The 2 groups consisted of XF media without glucose, and LPS + XF media without glucose. After 30 minutes of incubation at 37°C with 5% CO2, 1*μ*L phagocytotic beads was added. After another hour of incubation 20*μ*L anti-CD14-PE antibody was added. After another 30 minutes of incubation the cells were washed with PBS and resuspended in 400*μ*L PBS then analyzed with the Attune Nxt flow cytometer to determine the mean fluorescence intensity (MFI, the average amount of beads phagocytosed) and percentage of beads (% gated) phagocytosed by CD14+CD16- monocytes.

### Cytokine Expression Quantification using qPCR

To test differences in gene expression of 4 cytokines (IL-1*β*, IL-6, TNF*α*, IL-10), qPCR was performed on monocyte lysates from the Seahorse assay as we have previously described^26^. Relative expression levels were calculated using the comparative CT method using *β*2 microglobulin (B2M) as a control. B2M was picked as a control as there is evidence suggesting it’s the most stable reference gene in LPS-stimulated monocytes^30^.

### Statistical Analysis

Statistical analysis was performed using R software (R v. 3.5.1). Categorical demographic data was analyzed by chi-square test (sex, race). Continuous demographic and anthropometric data (age, height, weight, body mass index) were analyzed by independent-samples t-test between young and older subjects. For metabolic parameters all data followed a normal distribution according to Shaprio-Wilk tests. However, only difference in OCR (max-min) between groups had equal variances according to Levene’s test. Therefore independent-samples t-test was only used to calculate difference in OCR for metabolic parameters. Due to the unequal variances for the remaining parameters (max OCR, min OCR, area under the curve (AUC) kinetic OCR response) Mann-Whitney U tests were performed to test for significance between groups. For cytokine gene expression data at least 1 group for all genes did not meet the criteria for approximating normal distribution according to Shaprio-Wilk test, although all genes displayed equal variance between groups according to Levene’s test. Therefore, Mann-Whitney U tests were performed to test for significance between groups. For phagocytosis, all data met criteria for normal distribution and equality of variances according to Shapiro-Wilk and Levene’s test. Between group differences were determined by 2×2 (Condition x Group) ANOVA. As is standard, a p value < 0.05 was considered significant. Reported results are mean ± SEM.

## Acknowledgements

The authors would like to acknowledge the participants in this study.

## Funding

The study was supported by an American Heart Association grant (18AIREA33960189), startup funds, and a School of Health Studies Faculty Research Grant to BP.

## Availability of data and materials

The datasets and analytical scripts generated and analyzed during the current study are available in the FigShare repository, `10.6084/m9.figshare.c.4499219`.

## Author contributions statement

BP conceived and designed the study. BP and JY collected and analyzed the data. JY drafted the manuscript. BP and JY revised the manuscript drafts. All authors read and approved the final manuscript.

## Ethics approval and consent to participate

All study activities were approved by the Institutional Review Board at the University of Memphis (protocol #4361). Subjects provided informed consent and were free to withdraw at any time.

## Competing Interests

The authors declare that they have no competing interests.

## References

1. Colby, S. L. & Ortman, J. M. Projections of the size and composition of the US population: 2014 to 2060. Curr. Popul. Reports P25–1143, DOI: P25-1143 (2015). 1011.1669v3.

2. Azhar, G. & Wei, J. Y. The Demographics of Aging and Its Impact on the Cardiovascular Health. Curr. Cardiovasc. Risk Reports 9, 13, DOI: 10.1007/s12170-015-0441-x (2015).

3. Kennedy, B.K. et al. Geroscience: Linking Aging to Chronic Disease. Cell 159, 709–713, DOI: 10.1016/j.cell.2014.10.039 (2014).

4. Chatterji, S., Byles, J., Cutler, D., Seeman, T. & Verdes, E. Health, functioning, and disability in older adults—present status and future implications. The Lancet 385, 563–575, DOI: 10.1016/S0140-6736(14)61462-8 (2015).

5. Jacob, M. E. et al. Can a Healthy Lifestyle Compress the Disabled Period in Older Adults? J. Am. Geriatr. Soc. 64, 1952–1961, DOI: 10.1111/jgs.14314 (2016).

6. Franceschi, C. & Campisi, J. Chronic Inflammation (Inflammaging) and Its Potential Contribution to Age-Associated Diseases. The Journals Gerontol. Ser. A: Biol. Sci. Med. Sci. 69, S4–S9, DOI: 10.1093/gerona/glu057 (2014).

7. Salvioli, S. et al. Inflamm-Aging, Cytokines and Aging: State of the Art, New Hypotheses on the Role of Mitochondria and New Perspectives from Systems Biology. Curr. Pharm. Des. 12, 3161–3171, DOI: 10.2174/138161206777947470 (2006).

8. Franceschi, C., Garagnani, P., Vitale, G., Capri, M. & Salvioli, S. Inflammaging and ‘Garb-aging’. Trends Endocrinol. & Metab. 28, 199–212, DOI: 10.1016/j.tem.2016.09.005 (2017).

9. Michaud, M. et al. Proinflammatory Cytokines, Aging, and Age-Related Diseases. J. Am. Med. Dir. Assoc. 14, 877–882, DOI: 10.1016/j.jamda.2013.05.009 (2013).

10. López-Otín, C., Blasco, M. A., Partridge, L., Serrano, M. & Kroemer, G. The Hallmarks of Aging. Cell 153, 1194–1217, DOI: 10.1016/j.cell.2013.05.039 (2013). NIHMS150003.

11. Serbina, N. V., Jia, T., Hohl, T. M. & Pamer, E. G. Monocyte-Mediated Defense Against Microbial Pathogens. Annu. Rev. Immunol. 26, 421–452, DOI: 10.1146/annurev.immunol.26.021607.090326 (2008).

12. Albright, J. M. et al. Advanced Age Alters Monocyte and Macrophage Responses. Antioxidants & Redox Signal. 25, 805–815, DOI: 10.1089/ars.2016.6691 (2016).

13. Pence, B. D. & Yarbro, J. R. Aging impairs mitochondrial respiratory capacity in classical monocytes. Exp. Gerontol. 108, 112–117, DOI: 10.1016/j.exger.2018.04.008 (2018).

14. Raulien, N. et al. Fatty Acid Oxidation Compensates for Lipopolysaccharide-Induced Warburg Effect in Glucose-Deprived Monocytes. Front. Immunol. 8, DOI: 10.3389/fimmu.2017.00609 (2017).

15. Marsin, A.-S., Bouzin, C., Bertrand, L. & Hue, L. The Stimulation of Glycolysis by Hypoxia in Activated Monocytes Is Mediated by AMP-activated Protein Kinase and Inducible 6-Phosphofructo-2-kinase. J. Biol. Chem. 277, 30778–30783, DOI: 10.1074/jbc.M205213200 (2002).

16. Rodriguez-Prados, J.-C. et al. Substrate Fate in Activated Macrophages: A Comparison between Innate, Classic, and Alternative Activation. The J. Immunol. 185, 605–614, DOI: 10.4049/jimmunol.0901698 (2010).

17. Krawczyk, C. M. et al. Toll-like receptor-induced changes in glycolytic metabolism regulate dendritic cell activation. Blood 115, 4742–4749, DOI: 10.1182/blood-2009-10-249540 (2010).

18. O’Neill, L. A. J., Kishton, R. J. & Rathmell, J. A guide to immunometabolism for immunologists. Nat. Rev. Immunol. 16, 553–565, DOI: 10.1038/nri.2016.70 (2016).

19. Ravi, S., Mitchell, T., Kramer, P. A., Chacko, B. & Darley-Usmar, V. M. Mitochondria in monocytes and macrophages-implications for translational and basic research. The Int. J. Biochem. & Cell Biol. 53, 202–207, DOI: 10.1016/j.biocel.2014.05.019 (2014). NIHMS150003.

20. Vats, D. et al. Oxidative metabolism and PGC-1*β* attenuate macrophage-mediated inflammation. Cell Metab. 4, 13–24, DOI: 10.1016/j.cmet.2006.05.011 (2006).

21. Payne, B. A. & Chinnery, P. F. Mitochondrial dysfunction in aging: Much progress but many unresolved questions. Biochimica et Biophys. Acta (BBA) - Bioenerg. 1847, 1347–1353, DOI: 10.1016/j.bbabio.2015.05.022 (2015).

22. Houtkooper, R. H. et al. The metabolic footprint of aging in mice. Sci. Reports 1, 134, DOI: 10.1038/srep00134 (2011).

23. Balbir-Gurman, A., Yigla, M., Nahir, A. M. & Braun-Moscovici, Y. Rheumatoid Pleural Effusion. Semin. Arthritis Rheum. 35, 368–378, DOI: 10.1016/j.semarthrit.2006.03.002 (2006).

24. Chow, E. & Troy, S. B. The Differential Diagnosis of Hypoglycorrhachia in Adult Patients. The Am. J. Med. Sci. 348, 186–190, DOI: 10.1097/MAJ.0000000000000217 (2014).

25. Pence, B. D. Monocytes from older adults maintain capacity for metabolic compensation during glucose deprivation and lipopolysaccharide stimulation. figshare https://doi.org/10.6084/m9.figshare.c.4499219 (2019).

26. Pence, B. D. & Yarbro, J. R. Classical monocytes maintain ex vivo glycolytic metabolism and early but not later inflammatory responses in older adults. Immun. & Ageing 16, 3, DOI: 10.1186/s12979-019-0143-1 (2019).

27. Ong, S.-M. et al. The pro-inflammatory phenotype of the human non-classical monocyte subset is attributed to senescence. Cell Death & Dis. 9, 266, DOI: 10.1038/s41419-018-0327-1 (2018).

28. Aguilar-Ruiz, S. R. et al. Human CD16 + and CD16 - monocyte subsets display unique effector properties in inflammatory conditions in vivo. J. Leukoc. Biol. 90, 1119–1131, DOI: 10.1189/jlb.0111022 (2011).

29. Lachmandas, E. et al. Microbial stimulation of different Toll-like receptor signalling pathways induces diverse metabolic programmes in human monocytes. Nat. Microbiol. 2, 16246, DOI: 10.1038/nmicrobiol.2016.246 (2017).

30. Piehler, A. P., Grimholt, R. M., Ovstebo, R. & Berg, J. P. Gene expression results in lipopolysaccharide-stimulated monocytes depend significantly on the choice of reference genes. BMC Immunol. 11, 21, DOI: 10.1186/1471-2172-11-21 (2010).

